# A therapeutic vulnerability linking MNK1/2 inhibition and G1/S cyclin-dependent kinase blockade in triple-negative breast cancer

**DOI:** 10.64898/2026.09.11.750258

**Authors:** Qiyun Deng, Mehdi Amiri, Andia Moshari, Binadi Diddeniya, Parastoo Akbarnia, Ana A. Piric, Erika Prando Munhoz, Yasaman Bagherian, Zilan Li, Michael A. Bellucci, Niaz Mahmood, Michael Pollak, Sidong Huang, Mehdi Amiri, Nahum Sonenberg

**Affiliations:** Department of Biochemistry, McGill University, Montreal, Canada; Rosalind and Morris Goodman Cancer Institute, McGill University, Montreal, Canada; Lady Davis Research Institute, Jewish General Hospital, McGill University, Montreal, Canada

**Keywords:** Triple-negative breast cancer, functional genomics, translatome analyses, mRNA translational regulation, cell cycle regulation

## Abstract

Triple-negative breast cancer (TNBC) is a clinically challenging disease with limited therapeutic options. MAP kinase-interacting kinases 1 and 2 (MNK1/2)-mediated phosphorylation of eukaryotic initiation factor 4E (eIF4E) promotes oncogenic translation and represents a potential therapeutic target. We previously showed that loss of eIF4E phosphorylation suppresses metastasis but not primary tumor growth. Here, an shRNA screen surveying druggable genes unveiled cyclin-dependent kinase 4 (CDK4) as a genetic vulnerability to MNK1/2 inhibition in MDA-MB-231 TNBC cells. Although CDK4/6 inhibitors are approved for hormone receptor (HR)-positive, HER2-negative breast cancer, they are not approved for TNBC. We demonstrate that MNK1/2 inhibition synergizes with CDK4/6 blockade suppresses TNBC cell growth, a synergy also observed in another independent TNBC cell line (SUM159) using a broader G1/S CDK (CDK2/4/6) inhibitor. Integrated RNA sequencing and ribosome profiling revealed combination-specific changes in translation associated with mitotic checkpoint control and DNA repair. Our findings document MNK1/2 inhibition as a strategy to sensitize TNBC to G1/S CDK inhibition and provide a new avenue for therapeutic combination.

## Introduction

Triple-negative breast cancer (TNBC) is an aggressive breast cancer subtype that lacks estrogen receptor (ER), progesterone receptor (PR), and human epidermal growth factor receptor 2 (HER2) expression, with limited targeted therapeutic options^1^. CDK4/6 inhibitors, now a cornerstone of therapy for HR-positive, HER2-negative breast cancer, are not approved for TNBC^2^.

mRNA translation is frequently dysregulated in cancer to promote uncontrolled proliferation, survival, and metastatic progression^3,4^. Translation initiation of the vast majority of cellular mRNAs commences with the recognition of the 5’ cap structure (m7GpppN, where ‘N’ denotes any nucleotide and ‘m’ denotes a methyl group) by the eukaryotic translation initiation factor 4F (eIF4F) complex, which comprises the cap-binding protein eIF4E, the scaffolding protein eIF4G, and the RNA helicase eIF4A^4^. eIF4F promotes ribosome recruitment to the mRNA and plays a critical regulatory role in cap-dependent translation initiation^4^. As the limiting factor of the eIF4F complex^5^, eIF4E activity is tightly regulated through two major signaling pathways: sequestration by 4E-binding proteins (4E-BPs) downstream of the mechanistic target of rapamycin complex 1 (mTORC1), and phosphorylation at Ser209 by MAP kinase-interacting kinases 1 and 2 (MNK1/2)^6,7^. 4E-BPs act as translational repressors by competing with eIF4G for binding to eIF4E, thereby preventing assembly of the eIF4F complex^8^. In response to growth signals, mTORC1 phosphorylates 4E-BPs at multiple residues, reducing their binding to eIF4E, thus promoting mRNA translation initiation^9–11^. mTORC1 signaling promotes the translation of a subset of cancer-associated mRNAs^12–15^, whereas MNK1/2-mediated phosphorylation of eIF4E enhances the translation of a different subset of oncogenic mRNAs involved in tumor progression and metastasis^16–18^. Accordingly, elevated phosphorylated eIF4E (p-eIF4E) has been linked to poor prognosis across multiple cancer types^19–22^. MNK1/2 inhibition is effective in selected hematologic malignancies, including acute myeloid leukemia and diffuse large B-cell lymphoma^23,24^. In contrast, MNK1/2 inhibition drastically suppresses breast cancer metastasis, but not primary tumor growth in mice^16,18^, indicating that additional therapeutic vulnerabilities must be co-targeted to maximize its clinical potential. Several studies demonstrated that MNK1/2 inhibition enhances the potency of anticancer therapies in selected solid tumor models, including ER-positive breast cancer, colorectal cancer, and melanoma^17,25,26^. However, these studies were guided by existing mechanistic knowledge rather than systematic, unbiased functional screening.

We employed a synthetic lethal genetic screen followed by pharmacological validation and integrated translatomic and transcriptomic profiling to unveil genetic vulnerabilities associated with MNK1/2 inhibition in TNBC cells. We show that inhibition of the MNK1/2-eIF4E axis exacerbates cellular sensitivity to G1/S CDK blockade and describe the translational changes associated with this therapeutic combination.

## Results

### Inducible pooled shRNA screen identifies CDK4 as a potential synthetic lethal partner of MNK1/2

Abrogating p-eIF4E limits metastasis but not primary tumor growth in TNBC mouse models^16,18^. Leveraging synthetic lethality and functional genomics screening, we searched for combination therapies to target TNBC cell growth. The human TNBC cell line MDA-MB-231, which harbors a KRAS^G13D^ mutation that drives constitutive RAS-ERK-MNK1/2 signaling and most probably elevated eIF4E phosphorylation, was selected for an *in vitro* synthetic lethal screen^27^. The MNK1/2 inhibitor eFT508 was chosen to pharmacologically block eIF4E phosphorylation in the screen because of its high potency and selectivity towards MNK1 and MNK2^28^.

We employed a doxycycline-inducible short hairpin RNA (shRNA) library targeting the human druggable genome (hDGG) to increase the potential of clinical translation. The hDGG library covers 2263 therapeutically actionable genes, with five independent shRNAs per gene (**Supplementary Table 1**). At the experimental endpoint, the relative abundance of each shRNA construct was quantified by Illumina sequencing (**Fig. 1a**). The non-treated population (No Dox) served as a baseline control to assess library representation throughout the screening period. The doxycycline-induced, vehicle-treated (Dox + Vehicle) population served as a control for shRNAs that impaired cell fitness independently of drug treatment following shRNA induction. The doxycycline-induced, eFT508-treated (Dox + eFT508) population was used to identify shRNAs that were selectively depleted by MNK1/2 inhibition. Candidate targets were defined as shRNAs represented by at least 200 sequencing reads in the vehicle-treated population and depleted by at least 80% following eFT508 treatment (**Fig. 1b, Supplementary Table 2**). 277 shRNAs targeting 240 genes were identified (**Supplementary Table 3**). Gene Ontology Biological Processes analysis (see Methods) revealed an enrichment of genes involved in DNA replication, protein phosphorylation, and G2/M transition of the mitotic cell cycle (**Fig. 1c**). To prioritize candidates for validation, we chose the genes represented by at least two independent shRNAs that met our selection criteria and for which FDA-approved (**Table 1**) or direct inhibitors existed (**Supplementary Table 4**). Among these, two independent shRNAs targeting cyclin-dependent kinase 4 (*CDK4*) were depleted by more than 80% following eFT508 treatment (**Fig. 1b, d**). Since there are three CDK4/6 inhibitors approved in the clinic^2^, *CDK4* was selected for subsequent pharmacological validation.

**Figure 1.**
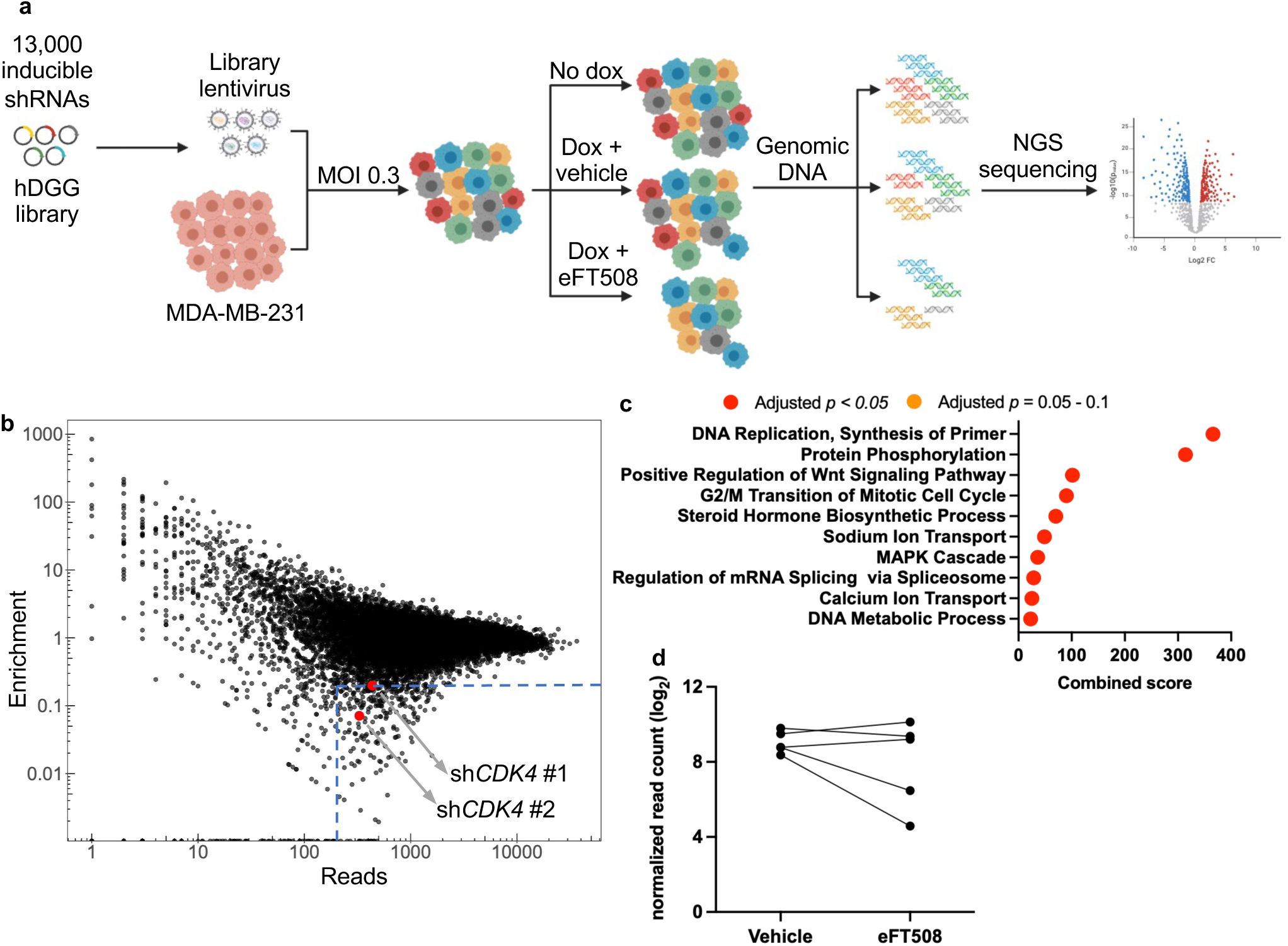
An inducible shRNA screen identifies CDK4 as a synthetic lethal candidate for p-eIF4E. (**a**) The schematic outline of the shRNA screen performed in MDA-MB-231 cells. Cells were transduced with the lentiviral hDGG library at an MOI of 0.3. Puromycin-selected cells were divided into three populations with different treatments as indicated. All three populations were cultured for eight doublings (11 days) to allow sufficient shRNA drop-out. Genomic DNA was extracted at the endpoint, and the inserted shRNA within the genome was PCR-amplified. The relative abundance of shRNAs in the hDGG library was analyzed by next-generation sequencing. (**b**) Visualization of the relative abundance of the shRNAs from the hDGG screen. The x-axis shows the number of reads in the vehicle population, and the y-axis shows the enrichment (the read ratio of eFT508 treated/vehicle). The arbitrary cutoff line (reads > 200, enrichment < 0.2) for significantly dropped-out shRNAs is indicated by the red dotted lines. Red dots indicate the two independent shRNAs targeting CDK4 that showed significant drop-out. (**c**) Gene Ontology Biological Process (GO-BP) enrichment analysis of the hDGG screen hits. shRNAs within the arbitrary cutoff line are considered for analysis. (**d**) log₂-normalized read counts of shRNAs targeting CDK4 in vehicle and eFT508-treated conditions at the end of the screen.

**Table 1:** Candidate genes identified in the shRNA screen with available FDA-approved inhibitors.

| Gene | Average Log<sub>2</sub> FC | FDA-approved inhibitors |
| --- | --- | --- |
| **CETP** | -3.6 | Torcetrapib |
| **TOP2A** | -3.5 | Etoposide, Teniposide, Doxorubicin, Idarubicin, Epirubicin, Mitoxantrone |
| **SCNN1D** | -3.5 | Amiloride |
| **AKR1B1** | -3.4 | Epalrestat |
| **CHRNA1** | -3.2 | Atracurium, Mivacurium, Pancuronium, Vecuronium |
| **CDK4** | -3.0 | Palbociclib, Ribociclib, Abemaciclib |
| **CACNG1** | -2.9 | Nimodipine |

### CDK4/6 inhibitors synergize with eFT508 to suppress MDA-MB-231 growth *in vitro*

We first evaluated palbociclib, which is the first-in-class FDA-approved CDK4/6 inhibitor for the treatment of HR-positive, HER2-negative breast cancer but is not approved for TNBC^29^. Although eFT508 alone had little effect on colony formation, combining it with palbociclib caused a significantly greater (20-30% depending on palbociclib dose) reduction in colony formation than palbociclib alone (**Fig. 2a, b**). To determine whether the enhanced growth inhibition reflected a synergistic interaction rather than an additive effect, drug interactions were quantified using the coefficient of drug interaction (CDI)^30,31^. CDI values between 0.7 and 1.0 indicated a synergistic interaction between eFT508 and palbociclib (**Fig. 2c**). Synergism was independently confirmed using the Highest Single Agent (HSA) model (**Fig. 2d**)^32^. Next, we evaluated ribociclib and abemaciclib, the other two FDA-approved CDK4/6 inhibitors, alone and in combination with eFT508 in MDA-MB-231 cells using real-time IncuCyte confluence-based proliferation assays. Despite differences in their potency and off-target inhibitory profiles^29,33,34^, all three CDK4/6 inhibitors exhibited synergistic antiproliferative effects when combined with eFT508 (**Fig. 2e-j**). Taken together, these findings demonstrate that pharmacological inhibition of CDK4/6 synergizes with MNK1/2 inhibition in suppressing the growth of MDA-MB-231 TNBC cells.

**Figure 2.**
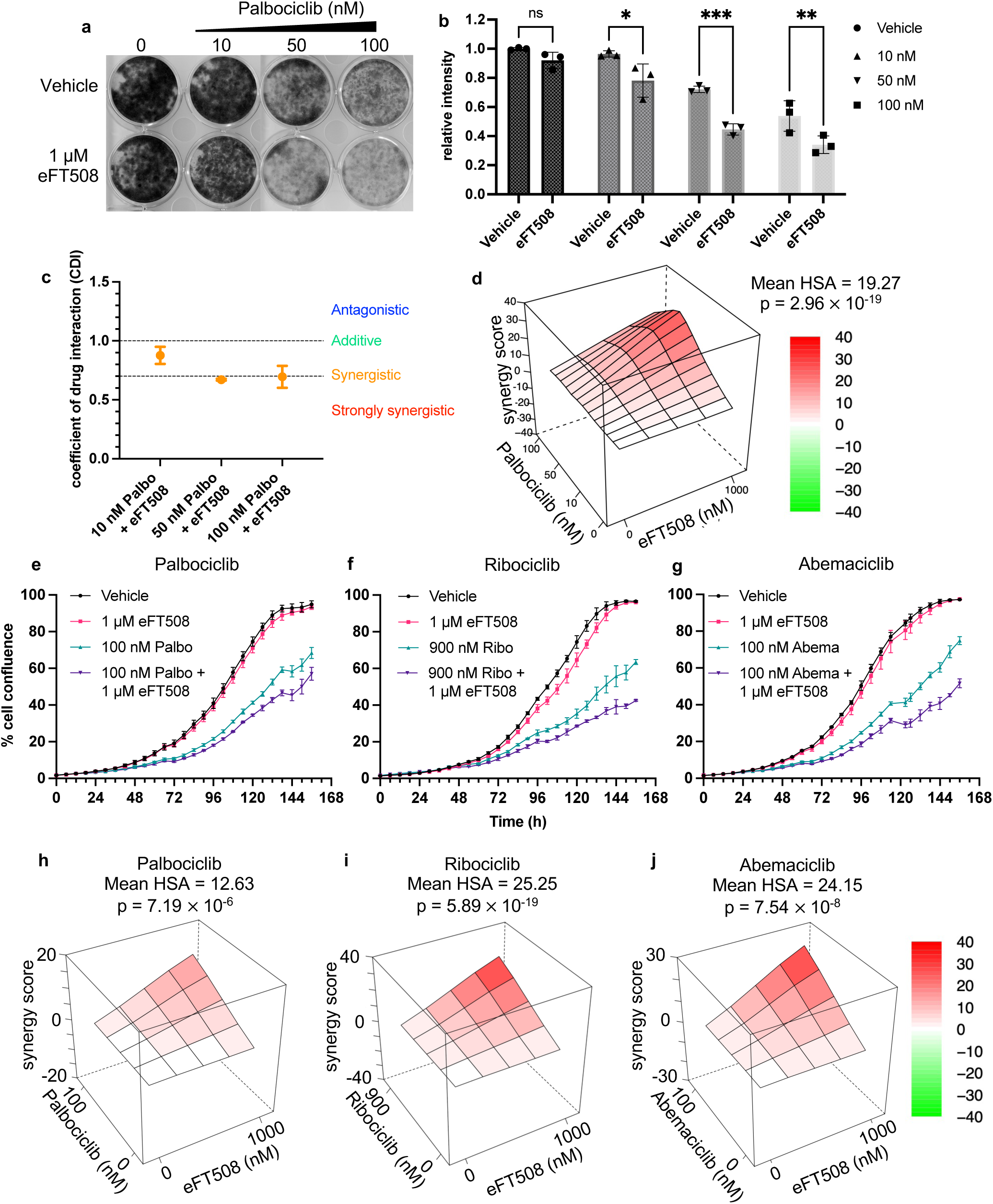
**Pharmacological CDK4/6 inhibition synergizes with MNK1/2 inhibition to inhibit TNBC growth *in vitro***. (**a**) Colony formation assay illustrating MDA-MB-231 cell growth upon different drug treatments. Cells were seeded in a 12-well plate and treated with different concentrations of palbociclib with or without 1 μM eFT508 for 8 days. For visualization, colonies were fixed using 4% paraformaldehyde and stained with crystal violet. (**b**) A bar graph showing the relative staining intensity of each well. The stained plates were scanned and quantified using the ImageJ ColonyArea plugin. *p < 0.0332, **p < 0.0021, ***p < 0.0002 (n = 3), two-way ANOVA test with correction for multiple comparisons using the Bonferroni test. Data presented as mean ± SD. (**c**) The coefficient of drug interaction (CDI) of the palbociclib-eFT508 combination in MDA-MB-231 cells. CDI > 1.0 indicates an antagonistic effect; CDI = 1.0 indicates an additive effect; CDI between 0.7 and 1.0 indicates synergy; and CDI < 0.7 indicates strong synergy. Error bars represent mean ± SD (n = 3). (**d**) HSA synergy scoring at multiple concentrations of palbociclib in combination with 1 μM eFT508 (n = 3). Drug combinations with synergy scores higher than 10, or lower than -10, were classified as synergistic or antagonistic interactions, respectively. (**e-g**) Representative real-time proliferation graphs of MDA-MB-231 cells treated with eFT508 in combination with (**e**) palbociclib, (**f**) ribociclib, and (**g**) abemaciclib. Images of the cells were taken every 6 hours using IncuCyte FLR, and cell proliferation was visualized based on the cell confluence in the images. (**h-j**) HSA synergy scoring for (**h**) palbociclib, (**i**) ribociclib, and (**j**) abemaciclib in combination with eFT508 (n = 3).

### CDK4/6 inhibition disrupts mTORC1 signaling and impairs eIF4E activity

To investigate the molecular basis underlying the synergism between CDK4/6 inhibition and eFT508, we examined key regulators of the G1/S cell cycle transition together with markers of mRNA translation control (**Fig. 3a**). As expected, all three CDK4/6 inhibitors markedly reduced phosphorylation of RB at Ser780, confirming effective inhibition of CDK4/6 activity. Total RB protein levels were also notably decreased, consistent with previous reports^35–37^. Cyclin D1 and CDK4 were upregulated by approximately 2-fold (**Fig. 3a, S1**), indicative of adaptive feedback following CDK4/6 inhibition^36,38,39^. As expected, eFT508 abolished phosphorylation of eIF4E, confirming inhibition of MNK1/2 activity (**Fig. 3a, S2**). Previous studies reported that CDK4/6 inhibition caused a reduction in the phosphorylation of 4E-BP1 ^40^, S6K and S6^41,42^. Consistent with these findings, CDK4/6 inhibitors alone reduced phosphorylation of 4E-BP1 (∼50 – 90%) (**Fig. 3a, S2**). Phosphorylation of both S6K and S6 was reduced (∼50 – 90%) following CDK4/6 inhibition, indicating suppression of mTORC1 signaling (**Fig.3b, S3**). These findings demonstrate that CDK4/6 inhibition, independent of MNK1/2-mediated eIF4E phosphorylation, also suppresses mTORC1-mediated translational control (4E-BP1, S6K, and S6), raising the possibility that this convergent effect on cap-dependent mRNA translation contributes to the synergy with eFT508.

**Figure 3.**
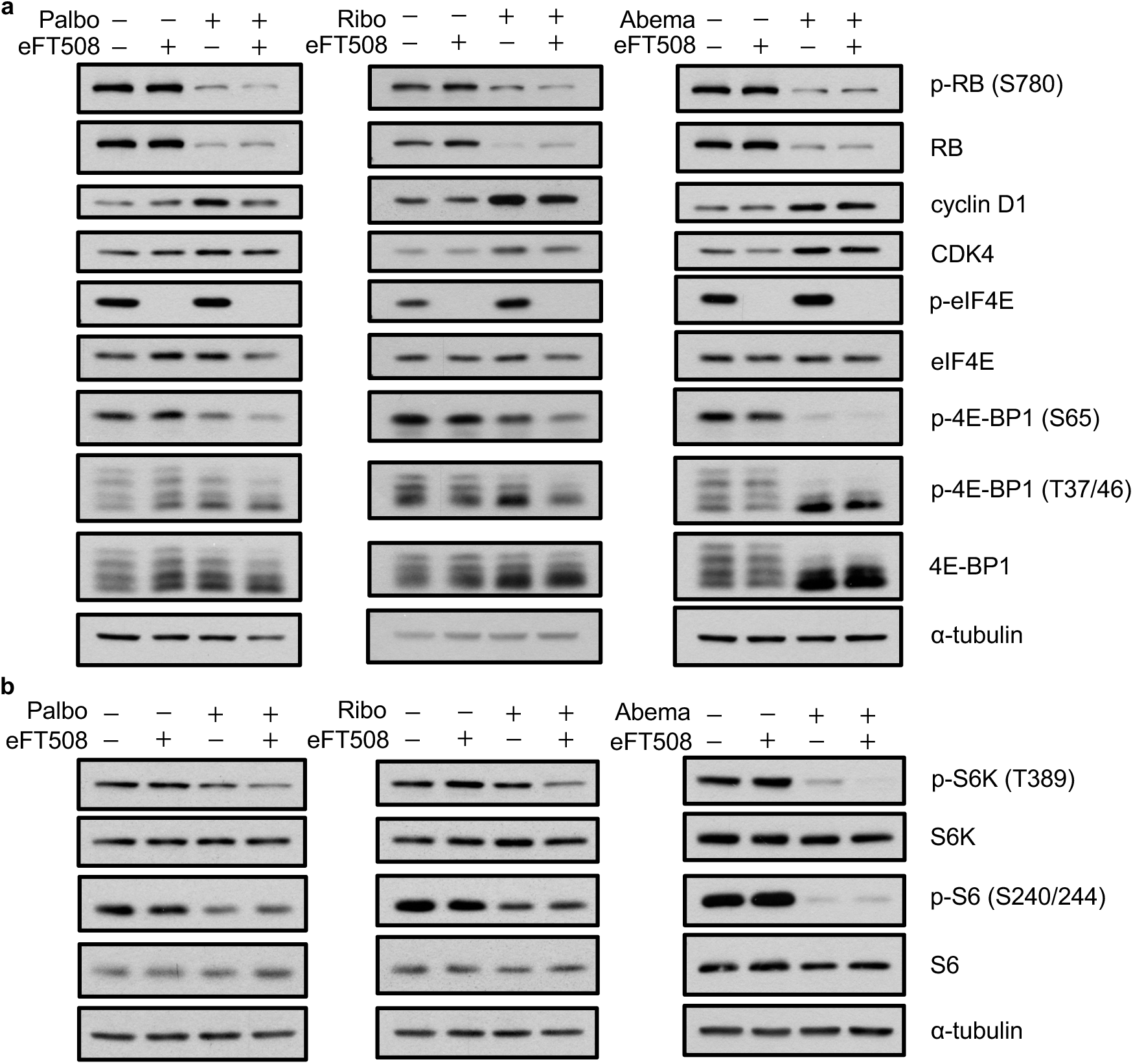
mTORC1 signaling is diminished when CDK4/6 is inhibited. (**a**) Representative western blot showing modulation of CDK4/6-RB pathway markers and 4E-BP1/eIF4E phosphorylation status upon treatment with 500 nM palbociclib, 2 μM ribociclib, or 500 nM abemaciclib, with or without 1 μM eFT508 for 24 hours (n = 3). (**b**) Representative western blot showing a reduction in phosphorylation of mTORC1 substrates when treated with different CDK4/6 inhibitors in the presence or absence of eFT508 (n = 3).

To determine whether pharmacological inhibition of mTOR signaling could explain the observed drug synergy, we examined two active-site mTOR inhibitors (AsTORi), INK128 and AZD8055, in combination with eFT508 in MDA-MB-231 cells using real-time proliferation assays. Both AsTORi inhibit mTORC1 and mTORC2 activity, thereby abolishing 4E-BP1 phosphorylation^43^. Neither inhibitor synergized with eFT508 to inhibit MDA-MB-231 cell growth despite efficient p-S6 and p-4E-BP1 reduction (**Fig. S4**). Thus, inhibition of the mTORC1 signaling alone is insufficient to account for the synergistic interaction between CDK4/6 inhibition and eFT508, indicating that other molecular mechanisms contribute to this synergy.

### Inhibition of G1/S cyclin-dependent kinases impairs mRNA translation

CDK4, CDK6, and CDK2 coordinately regulate G1/S cell cycle progression, as CDK4/6 initiate RB phosphorylation to promote E2F activation, and CDK2 reinforces this through a positive feedback loop^44^. Given that different cell lines exhibit varying dependencies on these kinases and that CDK2 activation confers resistance to CDK4/6 inhibitors, we next examined PF-06873600 (PF3600), a pan-CDK2/4/6 inhibitor, in targeting the G1/S checkpoint^45–48^. PF3600 synergized with eFT508 to suppress MDA-MB-231 cell growth to the same extent as CDK4/6 inhibitors (**Fig. 4a,c**). Importantly, the PF3600-eFT508 combination synergistically suppressed proliferation of another TNBC cell line, SUM159, established from a primary breast carcinoma^49^, demonstrating that the observed interaction extends to a genetically and phenotypically distinct TNBC model (**Fig. 4b,d**).

**Figure 4.**
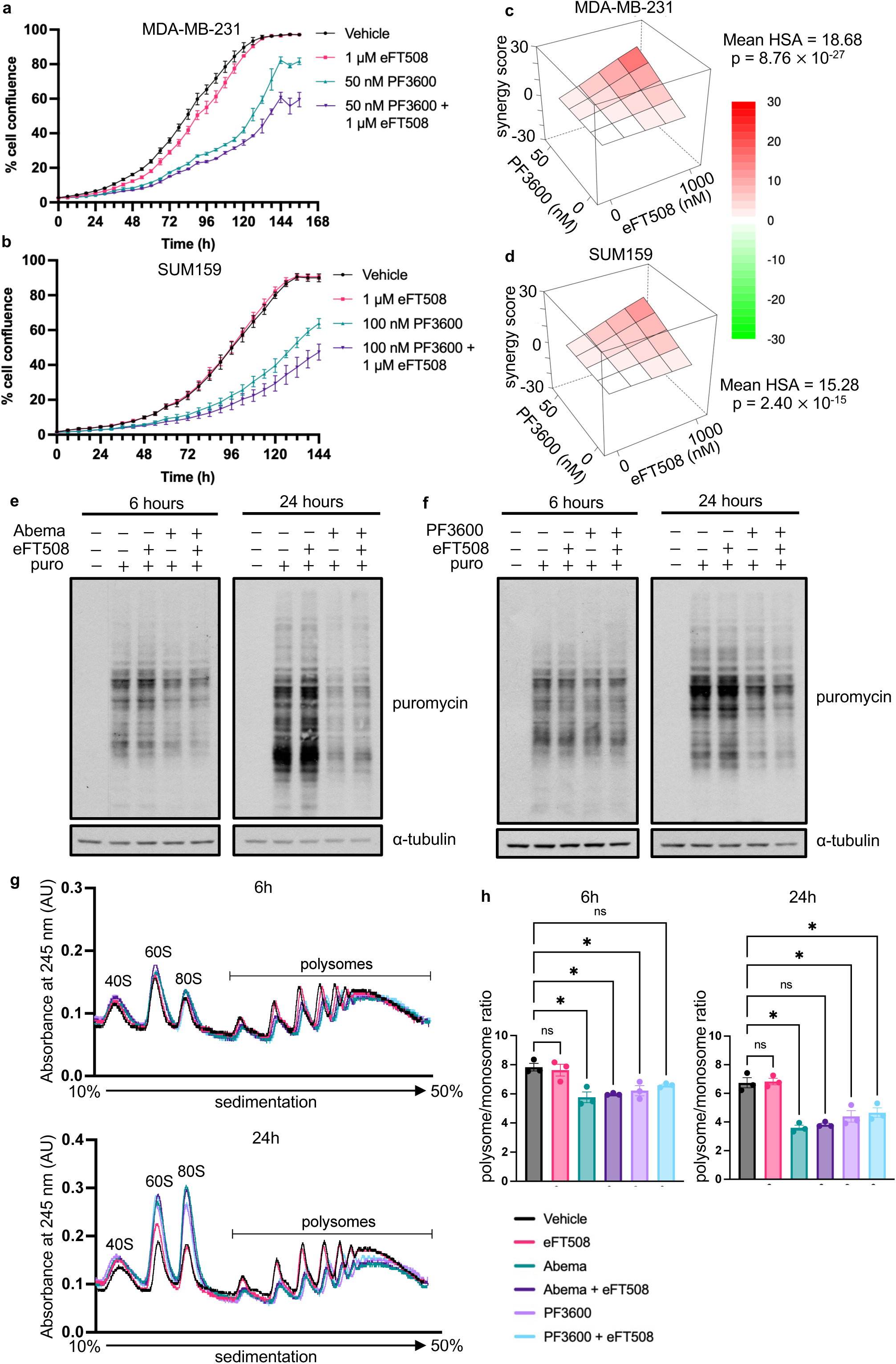
Inhibition of G1/S CDKs reduces general mRNA translation. (**a,b**) Representative real-time proliferation graphs of (**a**) MDA-MB-231 cells and (**b**) SUM159 cells treated with eFT508 in combination with PF3600. (**c,d**) HSA synergy scoring for eFT508 and PF3600 in (**c**) MDA-MB-231 cells and (**d**) SUM159 cells (n = 3). (**e,f**) Representative western blot showing puromycin incorporation in MDA-MB-231 cells upon treatment of (**e**) 500 nM abemaciclib and (**f**) 250 nM PF3600 with or without 1 μM eFT508. (**g**) Representative polysome profiles of MDA-MB-231 cells treated with 500 nM abemaciclib or 250 nM PF3600 with or without 1 μM eFT508 for 6 hours or 24 hours. (**h**) Quantification of polysome/monosome ratio of the polysome profiles, *p < 0.033, **p < 0.002 (n = 3), one-way ANOVA test with correction for multiple comparisons using the Dunnett test. Data presented as mean ± SD.

While all three CDK4/6 inhibitors similarly suppressed RB phosphorylation, abemaciclib caused a greater reduction in p-S6 phosphorylation (∼90%) than palbociclib or ribociclib (∼50%) (**Fig. 3**). We therefore selected abemaciclib, together with PF3600, for subsequent mechanistic studies, reasoning that the strongest pathway effect would maximize the chance of detecting combination-specific molecular changes. To determine whether the suppression of mTORC1 signaling was accompanied by reduced mRNA translation, we measured puromycin incorporation into nascent polypeptides following treatment with abemaciclib or PF3600, alone or in combination with eFT508, using Western blotting. Both CDK inhibitors reduced protein synthesis after 24 hours of treatment (∼30 – 70%), whereas eFT508 alone had no effect and the combination did not further reduce protein synthesis (**Fig. 4e,f**, **Fig. S5**). Consistent with these findings, polysome profiling revealed a time-dependent reduction in the polysome-to-monosome ratio following treatment with either abemaciclib or PF3600, with a ∼25% decrease at 6 hours and ∼50% at 24 hours (**Fig. 4g,h**). In contrast, eFT508 did not reduce polysome formation, consistent with earlier findings that MNK1/2-mediated eIF4E phosphorylation causes suppression of translation of only a small subset of selective mRNAs^16,50^.

### Combination-specific translational responses to eFT508 and G1/S CDK inhibition reveal candidate synergy mechanisms

Next, we investigated the effects of MNK1/2 inhibition on transcript-specific translational regulation using integrated translatome and transcriptome profiling. Since the G1/S CDK inhibitors abemaciclib and PF3600 suppressed mRNA translation in TNBC cells, we performed integrated ribosome profiling (Ribo-seq) and RNA sequencing (RNA-seq) to investigate the molecular mechanisms underlying the synergistic growth inhibition exerted by eFT508 and G1/S blockade. MDA-MB-231 cells were treated with vehicle, eFT508, abemaciclib, or their combination for 6 and 24 hours. Samples were subjected to parallel Ribo-seq and RNA-seq (**Fig. 5a**). Differential transcript abundance, ribosome occupancy, and translation efficiency (TE; defined as ribosome-protected fragment abundance normalized to mRNA abundance) were analyzed for each treatment condition. Sequencing library quality was assessed by examining ribosome footprint length distribution, read mapping characteristics, and metagene profiles, while reproducibility among biological replicates was evaluated by principal component analysis (PCA). Ribosome-protected fragments displayed the expected footprint length distribution with a peak centered at 36 nucleotides (nt) (**Fig. S6a**)^51^. Ribo-seq libraries exhibited an enrichment of reads mapping to coding sequences, and metagene analysis demonstrated the characteristic accumulation of ribosome footprints within coding regions (**Fig. S6b,c**). PCA demonstrated high reproducibility among biological replicates (**Fig. S6d**). To classify translationally regulated transcripts, RNA-seq and Ribo-seq log_2_ fold-changes were compared for each condition, and transcripts were categorized as forwarded (mRNA abundance change mirrored in ribosome occupancy without independent TE change), exclusive (TE changes without a corresponding change in mRNA abundance), intensified (mRNA abundance and TE changes act in the same direction, amplifying the net change in ribosome occupancy), and buffered (mRNA abundance change counteracted at the level of translation). Each category was further subdivided into upregulated and downregulated groups. Differential translation efficiency genes (DTEGs) were defined as transcripts with TE changes, which excludes forwarded transcripts.

**Figure 5.**
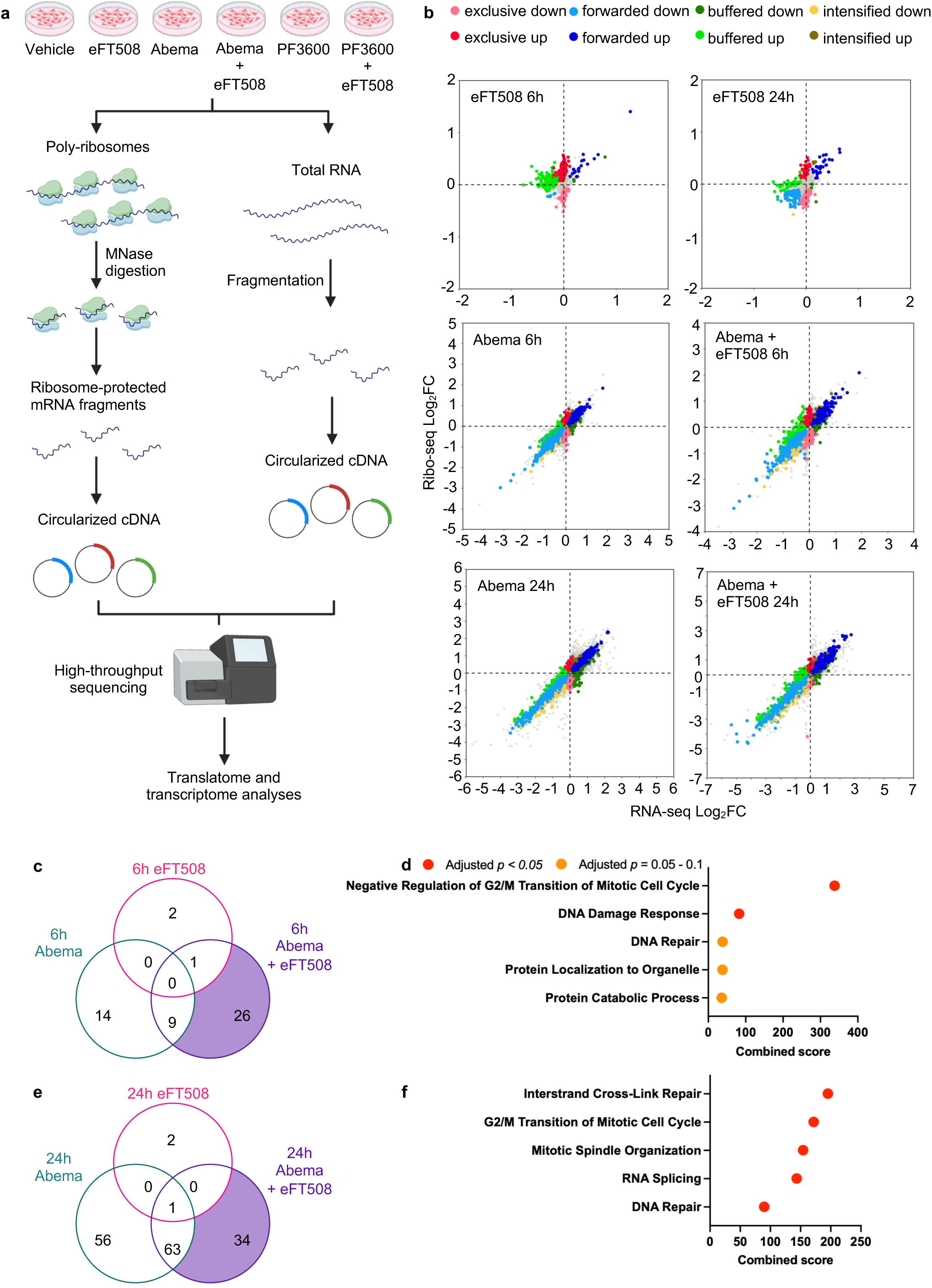
Integrated transcriptomic and translatomic analyses identify candidate pathways associated with the drug synergy. (**a**) Experimental design for RNA-seq and Ribo-seq of MDA-MB-231 cells treated with different drug combinations for 6 and 24 hours. Experiments were performed in biological triplicate. (**b**) Scatterplots showing the association between RNA-seq and Ribo-seq fold change log_2_FC between vehicle and different treatment groups. mRNAs with significant changes (adjusted p < 0.05 and log_2_FC > 0.2 or < -0.2) were classified based on coordinated changes in mRNA abundance, ribosome-protected fragments (RPFs), and translation efficiency (TE): intensified (concordant mRNA abundance and translational changes), forwarded (RPF changes proportional to mRNA abundance, with no TE change), exclusive (TE and RPF changes without changes in mRNA abundance), and buffered (opposing mRNA abundance and TE changes). (**c,e**) Venn diagrams showing the overlap of differential translation efficiency genes (DTGEs) following eFT508, abemaciclib, and combination treatment at (**c**) 6 hours and (**e**) 24 hours. Transcripts with adjusted p < 0.05 and TE log_2_FC < -0.5 were included. (**d,f**) Gene Ontology Biological Processes (GO-BP) enrichment analysis of the unique DTEGs in the eFT508 + abemaciclib group at (**d**) 6 hours and (**f**) 24 hours.

Integrated analysis of the transcriptome and translatome revealed distinct regulatory patterns across treatment groups and time points (**Fig. 5b, Supplementary Table 5**). eFT508 alone produced a modest change in TE at either time point: down-regulated DTEGs (log_2_FC < - 0.5, adjusted p-value < 0.05) include natural killer cell triggering receptor (NKTR), ADP ribosylation factor interacting protein 1 (ARFIP1) and breast cancer metastasis-suppressor 1-like protein (BRMS1L) at 6 hours; and S100 calcium binding protein A6 (S100A6), nucleobindin 2 (NUCB2), and programmed cell death 10 (PDCD10) at 24 hours. This demonstrates that eIF4E phosphorylation controls the translation of a small subset of mRNAs under basal unstressed conditions.

We reasoned that eIF4E phosphorylation may play an important role in the cellular response to stress^17^, such as G1/S CDK inhibition. We therefore asked whether combining eFT508 with G1/S CDK inhibitors produced a unique translational signature that explained the observed growth-suppressive synergy. Consistent with the established role of G1/S cyclin-dependent kinases in regulating E2F-dependent transcriptional programs^52^, abemaciclib predominantly affected mRNA abundance (**Fig. 5b**). To identify translational responses to combined MNK1/2 and abemaciclib inhibition, we compared down-regulated DTEGs unique to the eFT508-abemaciclib combination against those detected with either agent alone (**Fig. 5c,e**). At both 6 and 24 hours, combination-specific DTEGs were enriched for DNA repair and mitotic cell cycle progression. At 6 hours (# DTEGs = 26), the top terms were negative regulation of the G2/M transition, DNA damage response, and DNA repair (**Fig. 5d, Supplementary Table 5**). By 24 hours (# DTEGs = 34), DNA repair enrichment sharpened toward interstrand cross-link repair, alongside continued enrichment for the G2/M transition and mitotic spindle organization (**Fig. 5f, Supplementary Table 5**). This persistent enrichment suggests that combined MNK1/2 and CDK4/6 inhibition may compromise translation of DNA repair and mitotic regulators across both time points.

To determine whether these translational responses were shared across G1/S CDK inhibition more broadly, we profiled cells treated with eFT508 in combination with the CDK2/4/6 inhibitor, PF3600 (**Fig. S7a**). At 6 hours, the eFT508-PF3600 combination downregulated the TE of only 9 mRNAs, too few to support meaningful GO analysis (**Fig. S7b, Supplementary Table 5**). Notably, DNA cross-link repair 1A (DCLRE1A) and zinc finger GRF-type containing 1 (ZGRF1) are established DNA damage response genes^53,54^. Importantly, DCLRE1A was identified as a combination-specific intensified down DTEG in all three combinations: the 6-and 24-hour eFT508-abemaciclib combinations and the 6-hour eFT508-PF3600 combination. DCLRE1A functions in the DNA interstrand crosslink repair pathway and in the resolution of complex DNA double-strand breaks^54,55^. The translational downregulation of DCLRE1A across independent combination conditions provides strong support for impaired DNA repair capacity as a shared feature of both combinations. The eFT508-abemaciclib combination consistently generated larger and more robustly enriched DTEG sets than the eFT508-PF3600 combination; only 4 transcripts, chromosome 6 open reading frame 62 (C6ORF62), histone cluster 1 H2A family member K (HIST1H2AK), mitochondrially encoded NADH:ubiquinone oxidoreductase core subunit 4L (MT-ND4L), and nucleic acid binding protein 1 (NABP1), were uniquely downregulated by the eFT508-PF3600 combination at 24 hours (too few to support meaningful GO analysis).

Collectively, these translatome-level changes show that combined, but not single-agent, MNK1/2 and G1/S CDK inhibition selectively engenders reduced translation of mRNAs governing mitotic checkpoint control and DNA repair, thus informing about possible mechanisms of the synergistic growth suppression.

## Discussion

In this study, we report G1/S CDK blockade as a therapeutic vulnerability that can be exploited by MNK1/2 inhibition in TNBC. Using a pooled shRNA screen focusing on druggable genes, followed by pharmacological validation and integrated translatomic and transcriptomic profiling, we show that inhibition of G1/S cyclin-dependent kinases synergizes with the MNK1/2 inhibitor eFT508 to suppress TNBC cell growth. The drug synergy can be explained by translational suppression of mRNA encoding proteins involved in pathways governing mitotic progression and DNA repair. These findings provide a rationale for combining MNK1/2 inhibitors with G1/S-targeted therapies in TNBC.

Suppression of mTORC1 signaling and mRNA translation resulting from G1/S CDK inhibition is consistent with evidence linking G1/S cell cycle progression to mTORC1 regulation (**Fig. 3, 4e-h**)^42,56^. CDK4/6 phosphorylates tuberous sclerosis complex subunit 2 (TSC2) on Ser1452 and Ser1217, thereby dampening its inhibitory GAP function and promoting mTORC1 activation^42^. Accordingly, inhibition of CDK4/6 may restore TSC2-mediated suppression of mTORC1, consistent with the reduced phosphorylation of 4E-BP1, S6K, and S6 in our study. This conclusion is further supported by the observation that mTORC1 activity oscillates throughout the cell cycle, with its lowest activity in G1 and mitosis and highest during S and G2 phases^56^. This oscillation is mediated by the TSC complex independently of AKT or MEK/ERK and may also involve CDK2-dependent regulation of TSC^56^. These studies support a model in which G1/S CDKs coordinate cell-cycle progression through control of the TSC-mTORC1 axis, thereby coupling proliferation to protein synthesis. Despite the marked suppression of mTORC1 signaling, asTORi failed to phenocopy the synergy between eFT508 and G1/S blockade, indicating that inhibition of mTORC1 alone is insufficient to account for the therapeutic synergy in reducing TNBC cell growth. These findings suggest that additional downstream consequences of G1/S inhibition contribute to the enhanced response to MNK1/2 inhibition.

Although the genetic screen identified CDK4 as the top candidate synthetic lethal partner of MNK1/2 inhibition, our pharmacological studies suggested that the therapeutic interaction extends more broadly to the G1/S cyclin-dependent kinases. The absence of CDK2 and CDK6 from the screen results could be due to variable knockdown efficiencies, functional redundancy among closely related kinases, and cell-line-specific CDK dependencies^46,57,58^. CDK2, 4, and 6 compensate for one another to sustain G1/S progression, and activation of CDK2 is an important mechanism of resistance to CDK4/6 inhibitors^44–48^. These considerations led to our inclusion of the pan-CDK2/4/6 inhibitor PF3600, which demonstrated comparable synergy with eFT508 in MDA-MB-231 cells and in another TNBC model, SUM159 cells (**Fig. 4b**). These findings demonstrate that the synergy with MNK1/2 inhibition extends beyond CDK4 to the G1/S checkpoint more broadly, supporting combination strategies that target G1/S cyclin-dependent kinases as a class rather than CDK4 alone.

Despite causing a similar extent of inhibition in RB phosphorylation (**Fig. 3a**), the eFT508-abemaciclib combination exerted greater p-S6 reduction and combination-specific translational suppression than the eFT508-PF3600 combination (**Fig. 3b, S7**). Abemaciclib may have additional molecular responses beyond canonical CDK4/6 blockade, such as PIM1 inhibition^33,34^. Consequently, the enriched GO terms identified in the abemaciclib combination should be considered as candidate mechanisms requiring further experimental validation.

The antiproliferative effect of the drug combination is primarily driven by G1/S blockers (**Fig. 2, 4a-d**), whose therapeutic efficacy depends on an intact RB pathway^59^. Approximately 30% of TNBCs exhibit RB loss or inactivation and are therefore unlikely to benefit from this therapeutic approach^60,61^. To date, the best clinical success of CDK4/6 inhibitors is largely confined to combination with endocrine therapy for HR-positive, HER2-negative breast cancer^62,63^. Nevertheless, there is preclinical evidence demonstrating that a subset of TNBC cell lines, including MDA-MB-453 and CAL120, is sensitive to CDK4/6 inhibition^64^. This response variability may reflect the heterogeneous dependence of TNBC cells on individual G1/S CDKs. A recent study showed that cell lines exhibit distinct dependencies on CDK2, CDK4, and CDK6, with some TNBC cell lines relying predominantly on CDK2 rather than CDK4 or CDK6. For instance, MDA-MB-157 cells were sensitive to CDK2 depletion despite lacking dependence on CDK4 or CDK6 individually^46^. These findings suggest that selective CDK4/6 inhibition may fail to capture the full range of TNBC cells with therapeutically exploitable G1/S dependencies. Consistent with this, a phase II study of abemaciclib monotherapy (targeting CDK4/6 alone) in RB-positive metastatic TNBC did not meet its efficacy endpoint^65^. Broader inhibition of G1/S CDKs may therefore provide more comprehensive suppression of TNBC than CDK4/6 inhibition alone. Whether broader G1/S CDK inhibition can overcome this limitation remains unresolved. A clinical trial of the pan-CDK2/4/6 inhibitor PF-3600 enrolled patients with TNBC in the initial dose-finding cohort^66^. However, the subsequent efficacy-expansion cohorts evaluating antitumor activity did not enroll TNBC patients^66^. Further clinical development of PF-3600 was subsequently discontinued before its potential in TNBC could be specifically established, as Pfizer prioritized other CDK4- and CDK2-selective inhibitors^66^. Thus, although the heterogeneous CDK dependencies of TNBC provide a rationale for broader G1/S CDK blockade, achieving potent therapeutic activity may require rational combination strategies rather than monotherapies. In this context, combining G1/S CDK blockade with MNK1/2 inhibition may represent a strategy to enhance the anti-cancer activity and extend the therapeutic utility of CDK4/6 and CDK2/4/6 inhibitors in TNBC.

In the integrated translatome and transcriptome data, eFT508 treatment alone decreased the translation of a small subset of mRNAs. Several of these mRNAs, including NUCB2, S100A6, and PDCD10, encode factors that have documented pro-metastatic roles^67–69^. Translational suppression of these pro-metastatic factors by eFT508 is consistent with the anti- metastatic phenotype of eIF4E phosphorylation inhibition observed in breast cancer mouse models^16,18^. However, we found that the subset of mRNAs regulated by acute pharmacological MNK1/2 inhibition in MDA-MB-231 cells was distinct from the canonical p-eIF4E-dependent translational targets reported in previous studies, which were identified using genetic alteration of eIF4E itself (phospho-defective mutant)^16,17,50^. Direct comparison with the 35 mRNAs identified in wild-type relative to eIF4ES209A phospho-defective knock-in mouse embryonic fibroblasts (MEFs), including *Mmp3, Mmp9, Vegfc, Ccl2*, and *Pdgfra*^16^, showed no overlap with any mRNA translationally downregulated following eFT508 treatment. The differences may be due to the acute inhibition of MNK1/2 activity versus the absence of eIF4E phosphorylation of the Ser209Ala mutant, and/or the difference in cellular context between MEFs and the human TNBC cell line. Thus, the translational consequences of MNK1/2 inhibition are highly context- dependent and vary according to both tumor subtype and the method of MNK1/2 pathway perturbation (acute pharmacological inhibition versus chronic genetic manipulation).

The DTEGs selectively downregulated by combined MNK1/2 and G1/S CDK inhibition converged on two linked biological processes: G2/M checkpoint control and DNA repair. Notably, the DNA repair factor DCLRE1A was independently identified as an intensified down- target in our data set, exhibiting reduced translation efficiency on top of decreased mRNA abundance, in both the eFT508-abemaciclib and eFT508-PF3600 combinations. The convergence across two distinct G1/S CDK inhibitors likely reflects a genuine consequence of the combination therapy. Suppression of the DNA repair factors compromises repair of DNA lesions^70^. Disruption of centrosome and mitotic checkpoint-associated factors reduces accurate chromosome segregation^71,72^. Thus, combined MNK1/2 and G1/S CDK inhibition is expected to cause unresolved DNA damage and error-prone mitosis, which increases genome instability and the likelihood of a durable proliferative defect^73^. A related proteomic study combining MNK1/2 inhibition with palbociclib in melanoma and HR-positive breast cancer independently identified suppression of the same mitotic chromosome-segregation program, albeit through different effectors (survivin, AURKB, etc.)^26^. This consistency across distinct MNK1/2 inhibitors, cell contexts, and methods supports mitotic chromosome-segregation machinery as a shared consequence of combining MNK1/2 and CDK4/6 inhibition, which we extend here to TNBC for the first time. Future studies will be needed to determine whether these translational changes functionally contribute to the observed therapeutic synergy.

In summary, our study unveils G1/S CDK inhibition as a therapeutic vulnerability of MNK1/2 blockade in TNBC. These findings extend the clinical applicability of CDK4/6 inhibitors beyond their current use in HR-positive, HER2-negative breast cancer and provide a rationale for evaluating CDK2/4/6 inhibitors in this setting. By coupling unbiased functional screening with integrated translatomic and transcriptomic profiling, our study uncovers rational combination therapies and their candidate mechanisms more broadly. The convergence of both drug combinations on translational downregulation of DNA repair-related factors suggests a shared regulatory program worth pursuing mechanistically. Our findings provide a preclinical rationale for extending G1/S-targeted therapy to a subset of TNBC patients who currently lack this option.

## Materials and Methods Cell culture

The MDA-MB-231 cell line was a generous gift from Dr. Lynne-Marie Postovit (Queen’s University, Kingston, Canada), and SUM159 was from Dr. Jean Jacques Lebrun (McGill University, Montreal, Canada). MDA-MB-231 cells were cultured in Roswell Park Memorial Institute (RPMI) 1640 medium supplemented with 10% fetal bovine serum (FBS) and 1% penicillin/streptomycin. SUM159 was cultured in Ham’s F-12 medium supplemented with 5% FBS, 1% penicillin/streptomycin, 5μg/mL insulin, and 1μg/mL hydrocortisone. Cell lines were maintained at 37 °C and 5% CO_2_ for fewer than 20 passages. Cells were tested for mycoplasma contamination routinely via a PCR-based method and were authenticated by short-tandem-repeat (STR) profiling. Genomic DNA extracted from cell lines was analyzed by the Centre for Applied Genomics in Toronto, Canada.

## Compounds and antibodies

Tomivosertib (eFT508) was provided by eFFECTOR Therapeutics, Inc. (Solana Beach, CA, USA). Palbociclib (S1116), Abemaciclib (S5716), Ribociclib (S7440), and PF-06873600 (S8816) were purchased from Selleck Chemicals (Houston, TX, USA).

The antibodies used for Western blot are shown below in **Table 2**.

**Table 2.** Antibodies for Western Blotting.

| Targeted Protein | Company | Catalog number |
| --- | --- | --- |
| eIF4E | BD Biosciences | 610270 |
| p-eIF4E (S209) | Abcam | 76256 |
| RB | Cell Signaling Technology | 9313 |
| p-RB (S780) | Cell Signaling Technology | 9307 |
| α-Tubulin | Santa Cruz Biotechnology | 23948 |
| Cyclin D1 | Cell Signaling Technology | 2978 |
| Cyclin D3 | Abcam | 28283 |
| CDK4 | Abcam | 199728 |
| 4E-BP1 | Cell Signaling Technology | 9644 |
| p-4E-BP1 (T37/46) | Cell Signaling Technology | 2855 |
| p-4E-BP1 (S65) | Cell Signaling Technology | 9455 |
| S6K | Cell Signaling Technology | 9208 |
| p-S6K (T389) | Cell Signaling Technology | 9205 |
| S6 | Cell Signaling Technology | 2217 |
| p-S6 (S240/244) | Cell Signaling Technology | 2215 |
| puromycin | Millipore | MABE343 |

### Pooled synthetic lethal shRNA screen

shRNAs targeting human druggable genes (hDGG library) are in the lentiviral LT3GEPIR (pRRL) vector, where the GFP reporter and shRNA construct are under the doxycycline-inducible T3G promoter, and the puromycin resistance gene is under the PGK promoter acting as the selection marker. The library plasmid was obtained from the Genome Editing & Screening (GES) Core Facility at Memorial Sloan Kettering Cancer Center (MSKCC). The library lentivirus was generated using the protocol as described at https://portals.broadinstitute.org/gpp/public/resources/protocols.

MDA-MB-231 cells were transduced with the lentiviral library at MOI 0.3 and selected using 2μg/mL puromycin for 48 hours. Cells were pooled and plated at 3.25×10^6^ cells per 15-cm plate for three different conditions (no doxycycline control, doxycycline + vehicle, and doxycycline + eFT508). To ensure sufficient library representation, a 1000-fold coverage of the library size was maintained in each cell population. The medium was refreshed every two days, and cells were subcultured into new 15-cm plates every four days (maintaining the 1000-fold coverage). After 11 days of cell culture, genomic DNA was isolated from 1×10^7^ cells per condition using the High Pure PCR Template Preparation kit^74^.

shRNA constructs were extracted from 48 μg genomic DNA per condition by PCR amplification. The PCR amplification was done in two steps (PCR1 and PCR2) using the following cycling conditions: 98 °C, 30 s; (2) 98 °C, 10 s; 60 °C, 20 s; (4) 72 °C, 60 s; (5) to step (2), 16 cycles; (6) 72 °C, 5 min; (7) 4 °C. Indexes and adaptors for deep sequencing (Illumina) were added into PCR primers. 10 μL of PCR1 product was used as a PCR2 template for each condition. The final PCR2 products were purified using NucleoSpin Gel and PCR Clean-up kit (Takara Bio) according to the manufacturer’s manual. Subsequently, the purified PCR2 products were run on an 8% TBE-polyacrylamide gel to perform gel extraction as described in Ingolia’s paper^75^. The relative abundance of each shRNA construct was quantified by next-generation sequencing on the Illumina sequencing platform. The statistical analysis of the screen was processed using MAGeCK statistical software package (version 0.5.4)^76^.

The primers used are shown below in **Table 3**:

**Table 3.**
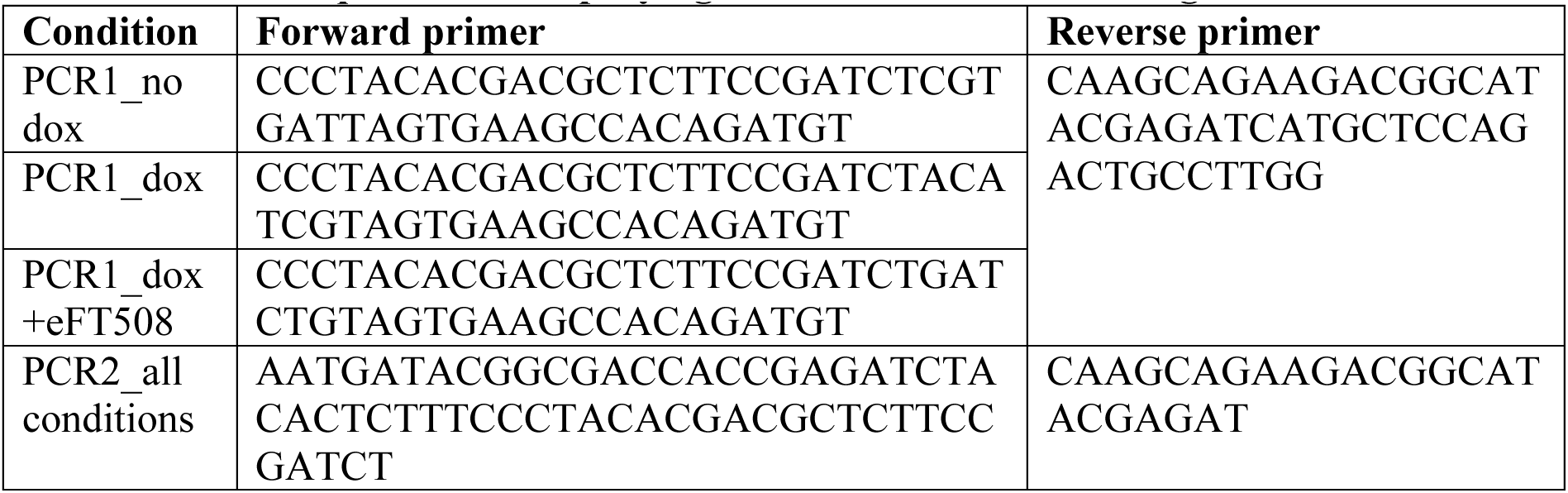
Primer sequence for amplifying shRNA inserts within the genomic DNA.

### GO-BP Enrichment Analysis

Gene Ontology Biological Process (GO-BP) enrichment analysis was performed using the Enrichr web-based gene set enrichment platform^77^. GO terms with adjusted *P*-value < 0.1 were considered significantly enriched. The top enriched GO-BP terms were visualized using dot plots generated in GraphPad Prism 11.

## Colony formation assay

5000 MDA-MB-231 cells were seeded into 12-well plates and cultured in the absence or presence of indicated drugs for 8 days. Drug-containing medium was replenished every two days. At the endpoint of the colony formation assay, cells were fixed with 4% paraformaldehyde in PBS and stained with crystal violet (0.5% w/v). Stained and dried plates were scanned at high resolution for ImageJ quantification using the ColonyArea plugin^78^. The ColonyArea plugin quantifies the stained colonies based on the colony size as well as its intensity, which can reflect the cell density difference^78^.

## IncuCyte cell proliferation assay

Cultured cells were seeded into 6-well plates at a density of 1 - 3 × 10^4^ cells per well. Cells were incubated overnight to allow attachment before drug treatment, refreshed every two days. Images of cells were taken using the phase-contrast setting every 6 hours using IncuCyte FLR or IncuCyte S3 machine depending on the cell types (Sartorius, Göttingen, Germany). Cell proliferation was quantified based on the cell confluence in the images.

## Coefficient of drug interaction (CDI) and highest single agent (HSA) synergy score calculation

Staining intensity data of colony formation assays were quantified using ImageJ. CDI for each palbociclib concentration was calculated using the equation: CDI = AB/(A × B) ^31^. “AB” is the relative staining intensity of the eFT508-palbociclib combination compared to the vehicle; “A” or “B” is the relative staining intensity of the groups treated with only one of the drugs compared to the vehicle. The HSA synergy scores were calculated using the SynergyFinder web application^79^. Drug combinations with synergy scores higher than 10, or lower than -10, were classified as synergistic or antagonistic interactions, respectively^32^.

## SDS-PAGE and Western blots

MDA-MB-231 cells were treated with corresponding inhibitors before protein extraction. Cells were lysed with RIPA buffer (50 mM Tris-HCl pH 7.4, 1% NP40, 0.1% SDS, 150 mM NaCl, 2 mM EDTA pH 8) supplemented with complete EDTA-free protease inhibitor cocktail (Roche) and phosphatase inhibitor cocktail 2 and 3 (Sigma-Aldrich). Lysates were incubated at 4 °C with rotation for 30 min and centrifuged at 15,000 × g for 15 min at 4 °C. Supernatant was collected and subjected to SDS-PAGE followed by Western blotting.

Proteins were denatured by adding 5× loading buffer (250 mM Tris-HCl, pH 6.8; 8 % w/v SDS; 0.2 % w/v bromophenol blue; 40 % v/v glycerol; 20 % v/v β-mercaptoethanol) and subjected to SDS-PAGE followed by transfer onto a 0.22 μm nitrocellulose membrane. Membranes were blocked in 5% skim milk solution (0.1 % TBST) and subjected to primary antibody incubation.

## Puromycin incorporation assay

MDA-MB-231 cells were seeded in 6-well plates and allowed to adhere overnight. Cells were treated with corresponding drugs for the indicated durations. Following treatment, cells were incubated with puromycin-containing medium (10 μg/mL) for 10 minutes, after which proteins were extracted using RIPA buffer.

## Polysome profiling

Polysome profiling was performed as described^51^. Briefly, MDA-MB-231 cells were treated with the indicated drugs before the CHX treatment (100 μg/mL) for 5 min at 37°C. Harvested cells were lysed in a hypotonic lysis buffer supplemented with 1.1% Triton X-100 and sodium deoxycholate. Cytoplasmic cell lysates were loaded on 10 - 50% sucrose gradients and fractionated after ultracentrifugation. Polysome-to-monosome ratios were calculated by integrating the area under the curve (AUC) of polysome and monosome peaks using OriginPro (OriginLab Corporation).

## Ribosome Profiling, RNA-sequencing, and Sequencing Data Alignment

Ribosome profiling was performed as described previously^51^. Briefly, MDA-MB-231 cells were treated with various inhibitors for 6 hours or 24 hours before harvest. Cells were collected in cycloheximide (CHX)-containing PBS and lysed in a 2X buffer containing 2% Triton X-100 and 2% sodium deoxycholate. MNase was used to digest the polysomes in the lysates (40.5U MNase per A260 Unit), then the digested polysomes were loaded on 10-50% sucrose gradients and separated using ultracentrifugation. The fractions containing the digested monosomes were collected using a collector and UV optical unit. These fractions were then subjected to RNA extraction by TRIzol (Invitrogen, Cat#15596018) and followed by library preparation as described previously^51^. For RNA-seq, total RNA was extracted from 50 uL of the undigested lysates using TRIzol. rRNA depletion was performed using the NEBNext rRNA Depletion Kit v2(Human/Mouse/Rat) on 1 ug of total RNA according to the manufacturer’s protocol. The depleted RNA samples were fragmented using NEBNext Magnesium RNA Fragmentation Module before processing to library preparation as described above. Sequenced libraries were trimmed, aligned, and statistically analyzed as described previously^51^.

## Acknowledgements

We thank Lynne-Marie Postovit (Queen’s University) for the MDA-MB-231 cell line and Jean Jacques Lebrun (McGill University) for the SUM159 cell line. shRNA screen sequencing was performed at the Donnelly Sequencing Centre (Toronto, Canada), and RNA-seq and Ribo-seq libraries were sequenced on an Illumina NovaSeq 6000 System at Novogene Inc. (Sacramento, CA, USA). The work was supported by grants from the Cancer Research Society (1051778, 1449911) and Canadian Institutes of Health Research (CIHR) (MBC-162042) to N.S., a Goodman Cancer Institute Canderel Studentship and a Fonds de recherche du Québec – Santé (FRQS) doctoral studentship to Q.D.

## Author contributions

N.S., M.A., and Q.D. conceived the project and designed the experiments. Q.D. performed most experiments and data analysis. A.M., B.D., and P.A. assisted in the polysome profiling and Ribo- seq. A.A.P. and Y.B. helped with western blotting and analysis. Z.L. contributed to the colony formation assays and IncuCyte proliferation assays. E.P.M. and M.A. conducted the shRNA screen sequencing analysis. M.A. conducted the Ribo-seq and RNA-seq analysis. M.A.B. contributed to data analysis and graphing. S.H. provided important guidance on the shRNA screen framework, techniques, and data interpretation. M.P. provided valuable clinical insight. Q.D., M.A., N.M., and N.S. interpreted the data and wrote the manuscript. All authors discussed the results and edited the manuscript.

## Resource availability

Further information and requests for resources should be directed to the lead contacts, Nahum Sonenberg and Mehdi Amiri. This paper does not report original code. Any information required to reanalyze the data reported in this work is available from the lead contacts upon request.

---

**Figure S1.**
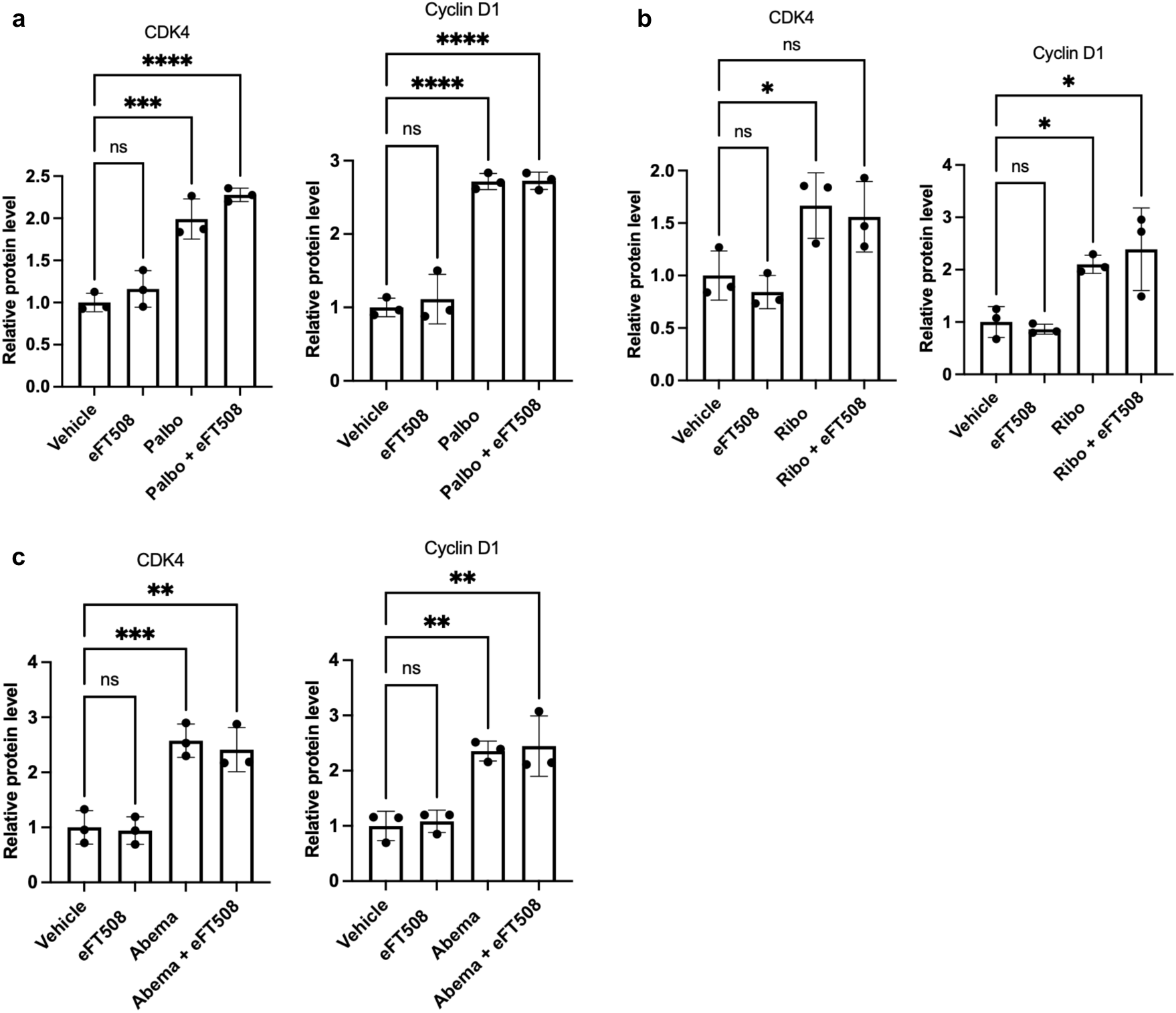
Statistical analysis of immunoblot quantification corresponding to. Fig 3a. (**a-c**) Quantification of CDK4 and cyclin D1 levels in MDA-MB-231 cells treated with eFT508 in combination with (**a**) palbociclib, (**b**) ribociclib and (**c**) abemaciclib. *p < 0.033, **p < 0.002, ***p < 0.0002, ****p < 0.0001 (n = 3), one-way ANOVA test with correction for multiple comparisons using the Dunnett test. Data presented as mean ± SD.

**Figure S2.**
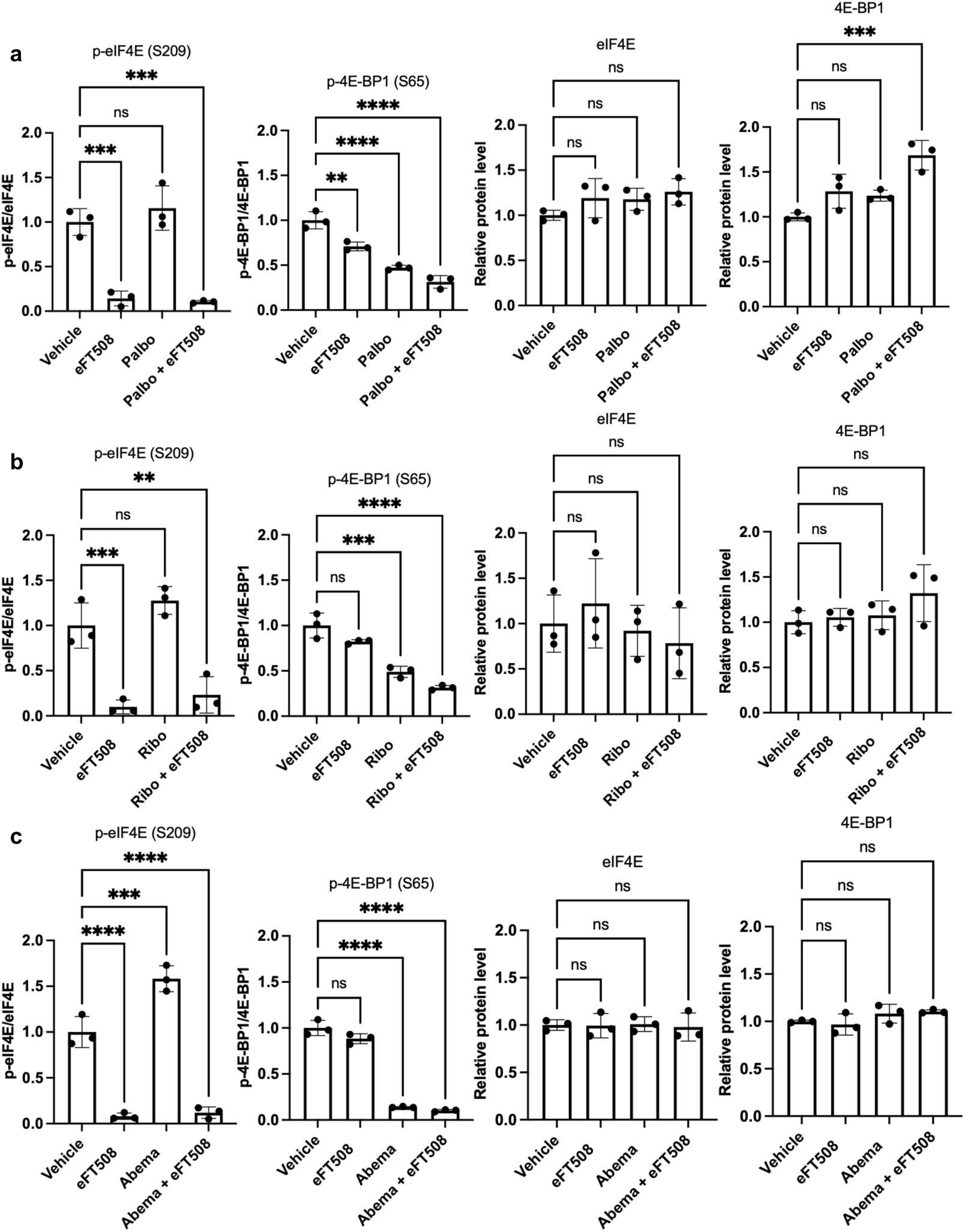
Statistical analysis of immunoblot quantification corresponding to. Fig 3a. (**a-c**) Quantification of p-eIF4E/eIF4E, p-4E-BP1 (S65)/4E-BP1, total eIF4E, and total 4E-BP1 of MDA-MB-231 cells treated with eFT508 in combination with (**a**) palbociclib, (**b**) ribociclib and (**c**) abemaciclib. *p < 0.033, **p < 0.002, ***p < 0.0002, ****p < 0.0001 (n = 3), one-way ANOVA test with correction for multiple comparisons using the Dunnett test. Data presented as mean ± SD.

**Figure S3.**
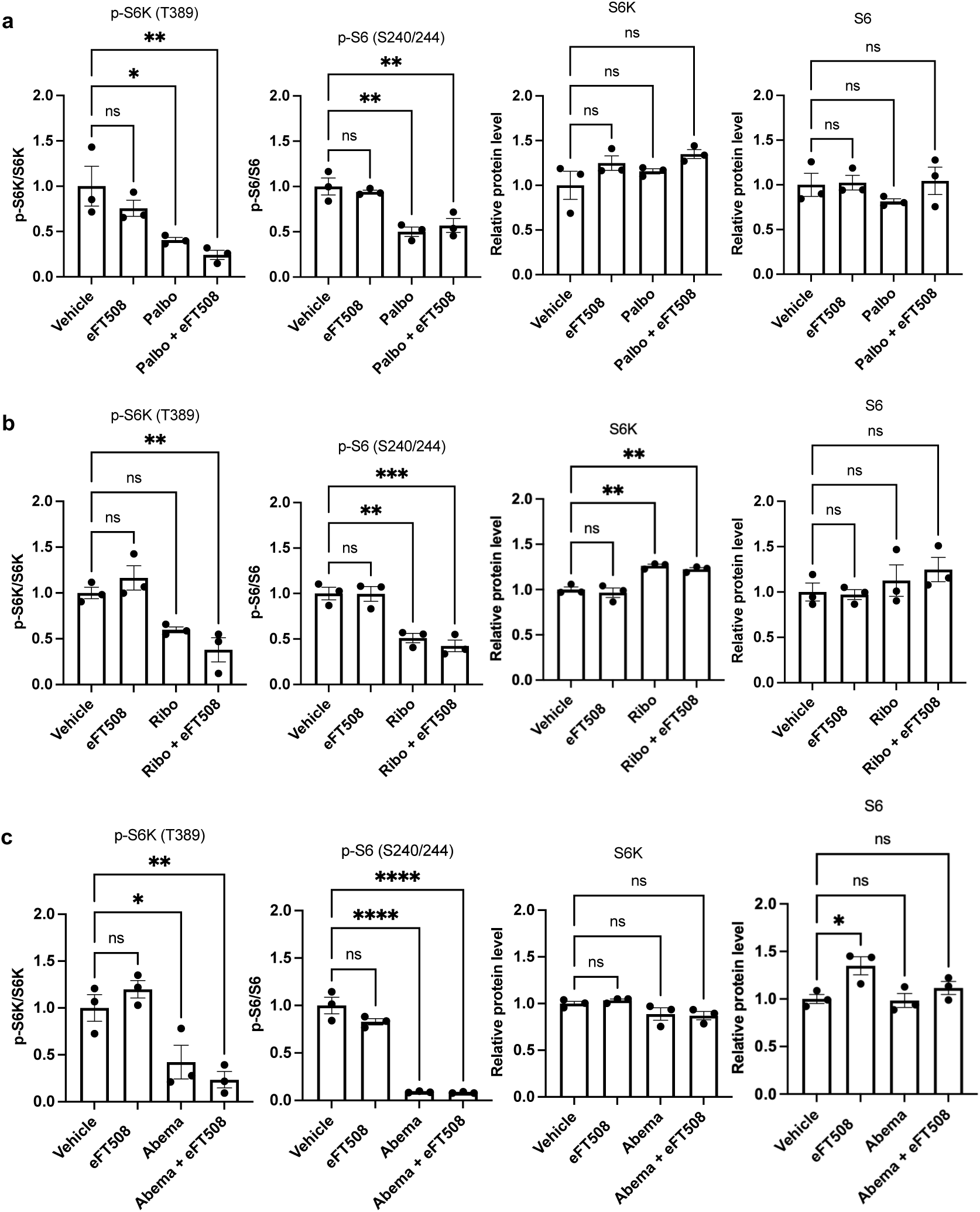
Statistical analysis of immunoblot quantification corresponding to. Fig 3b. (**a-c**) Quantification of p-S6K/S6K, p-S6/S6, total S6K, and total S6 of MDA-MB-231 cells treated with eFT508 in combination with (**a**) palbociclib, (**b**) ribociclib and (**c**) abemaciclib. *p < 0.033, **p < 0.002, ***p < 0.0002, ****p < 0.0001 (n = 3), one-way ANOVA test with correction for multiple comparisons using the Dunnett test. Data presented as mean ± SD.

**Figure S4.**
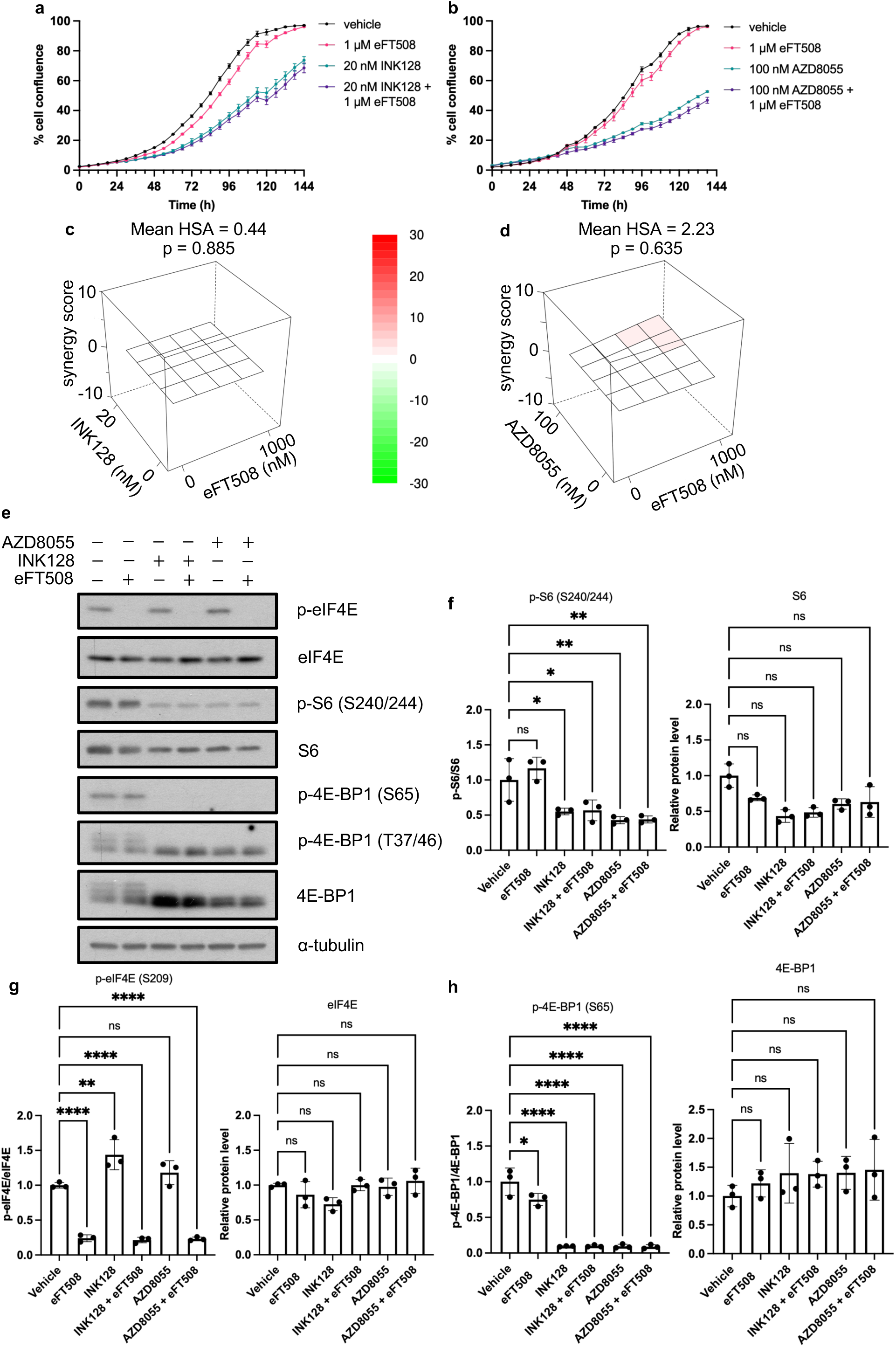
**Active-site mTOR inhibitors do not synergize with eFT508 in MDA-MB-231 cells**. Representative real-time proliferation graphs of MDA-MB-231 cells treated with eFT508 in combination with (**a**) INK128 or (**b**) AZD8055. (**c,d**) HSA synergy scoring for eFT508 and (**c**) INK128 or (**d**) AZD8055 in MDA-MB-231 cells (n = 3). (**e**) Representative western blot showing reduced phosphorylation of mTORC1 substrates after treatment with different AsTORi, with or without eFT508 (n = 3). (**f-h**) Quantification of (**f**) p-S6/S6, total S6, (**g**) p-eIF4E/eIF4E, total eIF4E, (**h**) p-4E-BP1, and total 4E-BP1 of MDA-MB-231 cells treated with eFT508 in combination with 20nM INK128 or 100nM AZD8055 for 24 hours. *p < 0.033, **p < 0.002, ****p < 0.0001 (n = 3), one-way ANOVA test with correction for multiple comparisons using the Dunnett test. Data presented as mean ± SD.

**Figure S5.**
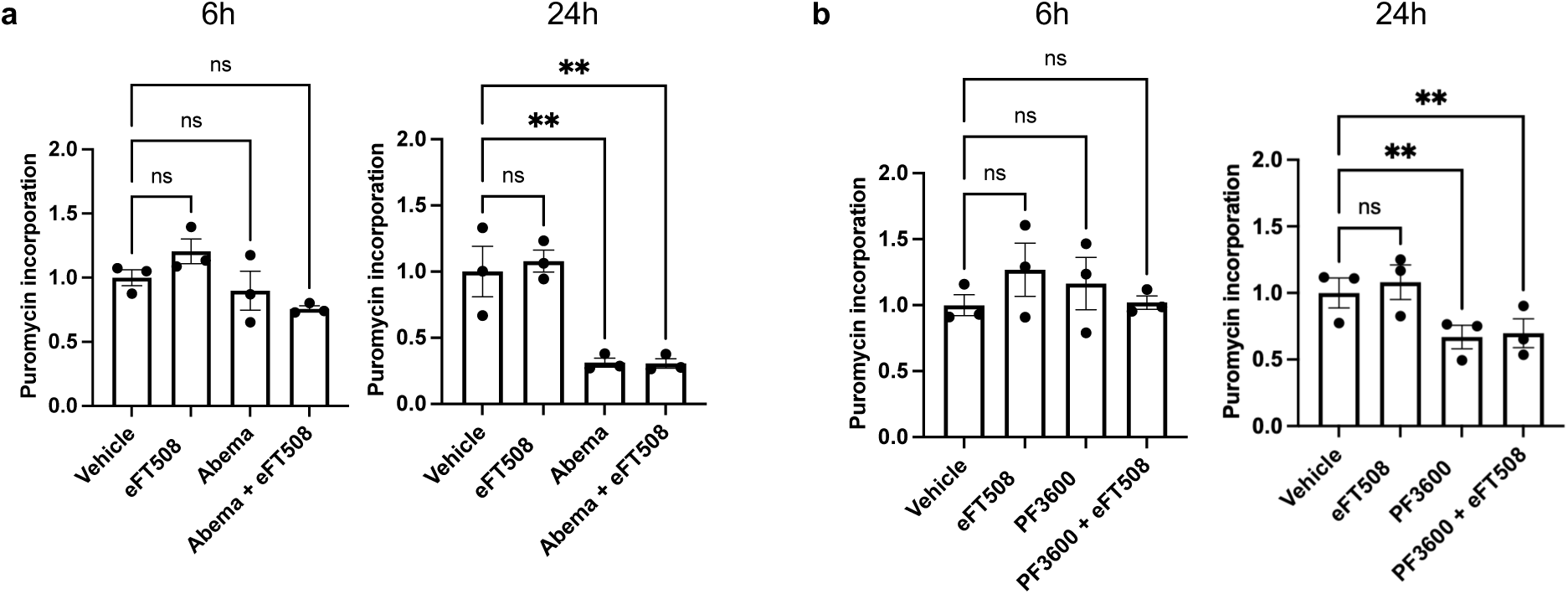
Statistical analysis of puromycin incorporation immunoblot quantification corresponding to. Fig 4e**,f**. Quantification of puromycin incorporation in MDA-MB-231 cells treated with eFT508 in combination with (**a**) abemaciclib or (**b**) PF3600, *p < 0.0332, **p < 0.0021 (n = 3); one-way ANOVA test with correction for multiple comparisons using the Dunnett test. Data presented as mean ± SD.

**Figure S6.**
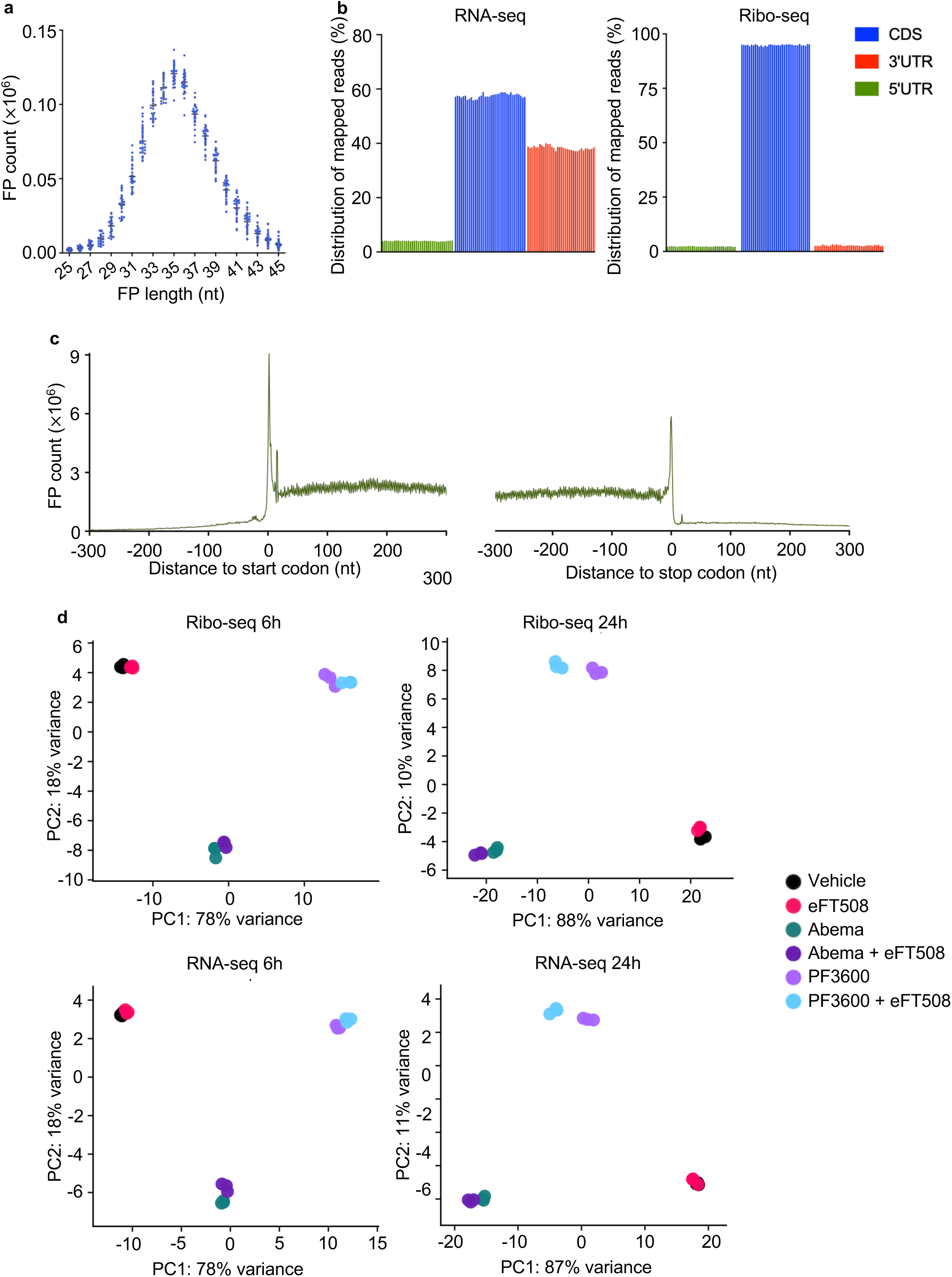
**Quality control metrics for Ribo-seq and RNA-seq libraries**. (**a**) Length distribution of ribosome-protected footprints (RPFs). (**b**) Percentage of RNA-seq and Ribo-seq reads mapping to the 5’ untranslated region (5’ UTR), coding sequence (CDS), and 3’ untranslated region (3’UTR). (**c**) Metagene analysis of Ribo-seq libraries showing the distribution of ribosome footprints aligned relative to the start and stop codon. (**d**) Principal component analysis (PCA) of RNA-seq and Ribo-seq datasets demonstrating reproducibility between biological replicates.

**Figure S7.**
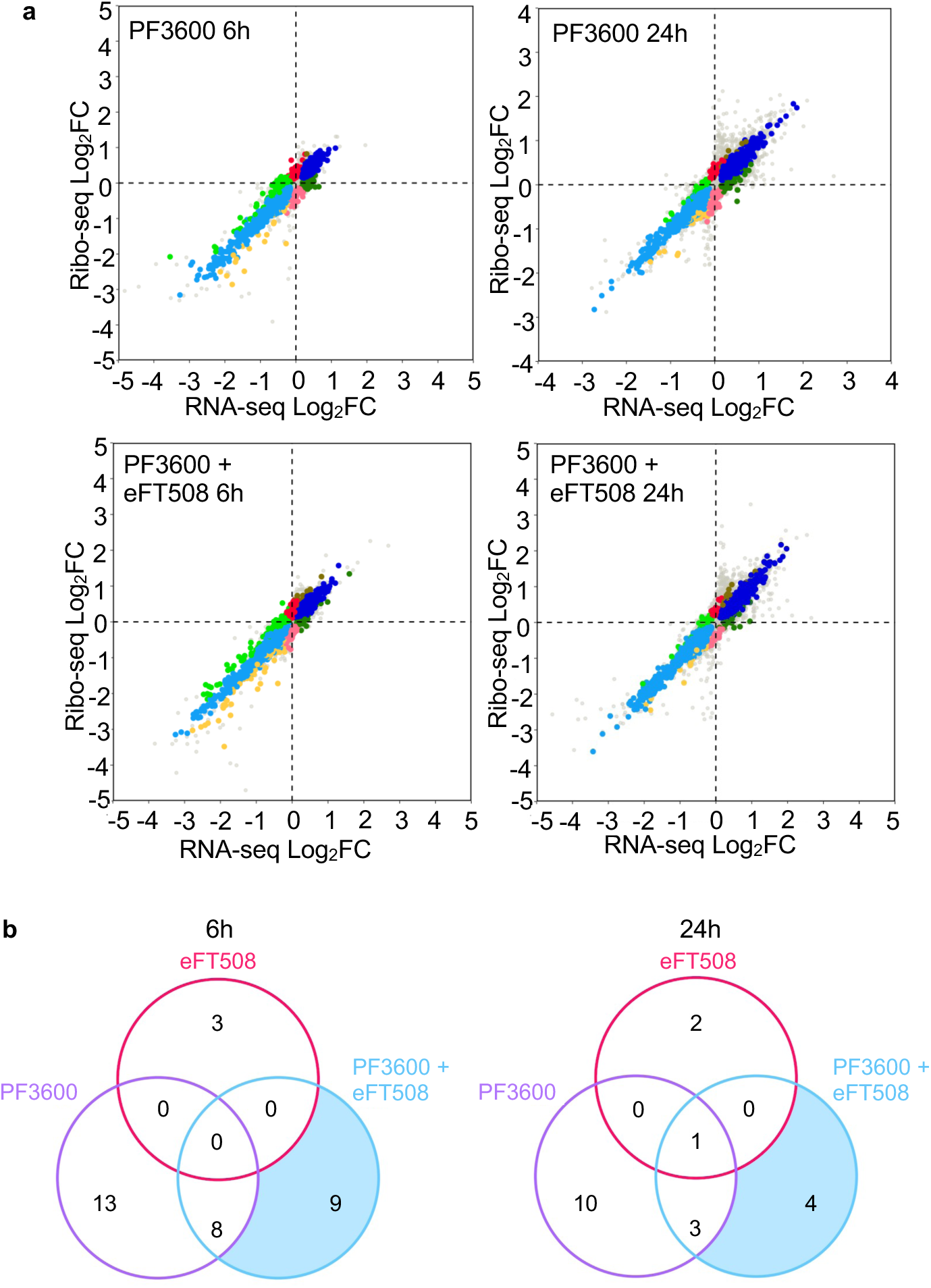
Integrated translatome and transcriptome profiling of PF3600-treated MDA- MB-231 cells with or without eFT508. (**a**) Scatterplots showing the relationship between RNA- seq and Ribo-seq log₂ fold changes (log₂FC) following treatment with PF3600 alone or in combination with eFT508 for 6 and 24 hours. Transcripts were classified as described in Fig. 5. (**b**) Venn diagrams showing the overlap of DTEGs following eFT508, PF3600, and combination treatment at 6 and 24 hours. Transcripts with adjusted p < 0.05 and TE log_2_FC < -0.5 were included.

